# Myelin-reactive B cells exacerbate the severity of CD4^+^ T cell-driven CNS autoimmunity in an IL-23-dependent manner

**DOI:** 10.1101/721696

**Authors:** Mohamed Reda Fazazi, Prenitha Mercy Ignatius Arokia Doss, Resel Pereira, Neva Fudge, Aryan Regmi, Charles Joly-Beauparlant, Irshad Akbar, Asmita Pradeep Yeola, Benoit Mailhot, Joanie Baillargeon, Philippe Grenier, Nicolas Bertrand, Steve Lacroix, Arnaud Droit, Craig S. Moore, Olga L. Rojas, Manu Rangachari

## Abstract

Multiple sclerosis (MS) is an autoimmune disorder of the central nervous system (CNS) that has traditionally been considered a T cell-mediated disease. However, accumulating evidence points to a crucial role for B cells in disease processes. IgH^[MOG]^ mice possess a transgenic immunoglobulin heavy chain derived from a monoclonal antibody specific to myelin oligodendrocyte glycoprotein (MOG), a key target for autoimmune responses. Here, using the experimental autoimmune encephalomyelitis (EAE) model of MS, we investigated the susceptibility of IgH^[MOG]^ mice to CD4^+^ T cell-driven disease that was induced by active immunization with MOG_[35-55]_ autoantigen. We found that immunized IgH^[MOG]^ mice rapidly developed severe EAE, characterized by the aggregation of T and B cells in the CNS meninges. We observed an increased presence of class-switched and inflammatory cytokine-positive B cells in the IgH^[MOG]^ CNS, as well as a greater frequency of IL-17- and GM-CSF-producing CD4^+^ T cells. Production of the Th17 maintenance factor IL-23 was increased from IgH^[MOG]^ CNS-infiltrating B cells, and *in vivo* blockade of IL-23p19 strongly attenuated disease severity in IgH^[MOG]^ mice. Strikingly, we observed an increased frequency of PD-1^+^CXCR5^-^ T peripheral helper (Tph)-like cells in the CNS parenchyma and dura mater of IgH^[MOG]^ mice. Both Tph accumulation in the CNS, as well as meningeal inflammation, were again sharply reduced upon IL-23p19 blockade *in vivo*. Notably, CNS-infiltrating B and Tph cells from IgH^[MOG]^ mice showed an upregulation of genes related to neurodegeneration and oxidative phosphorylation, and IL-23 blockade reduced ROS production from these cells *in vivo*. Altogether, these data show that MOG-specific B cells contribute to severe CD4^+^ T cell-driven EAE by promoting CNS accumulation of Th17 and Tph cells, as well as tertiary lymphoid organs in the CNS meninges, in an IL-23 dependent manner.

## Introduction

Multiple sclerosis (MS) is a chronic autoimmune disease in which the adaptive immune system launches an attack against central nervous system (CNS) proteins, such as myelin. MS affects more than 2 million people worldwide (1). Approximately 80% of patients present an initially relapsing-remitting (RR) disease course for which there are now more than 10 disease-modifying therapies available. However, 30-60% of these RR patients will eventually transition to a chronically worsening secondary progressive (SP) phase, for which treatment options are limited (2). Pathophysiological mechanisms in progressive MS are thus of intense current interest (3).

T cells, and CD4^+^ T cells in particular, have historically been the most intensively studied players in the immune pathogenesis of MS. However, it has become increasingly clear that B cells additionally play important roles in MS. Clonally expanded B cells are present in the cerebrospinal fluid (CSF) and MS plaques (4–6), and the presence of meningeal follicles adjacent to cortical lesions is associated with disease progression (7, 8). Further, antibodies against myelin oligodendrocyte glycoprotein (MOG), a key component of myelin, were found in active MS lesions (9). Crucially, the B cell-targeting anti-CD20 drugs rituximab(10), ocrelizumab (11) and ofatumumab (12) improve RRMS, and ocrelizumab is the only FDA approved drug for primary progressive MS (13). Curiously, though, antibody-secreting plasmablasts do not express CD20 (14). This suggests that the pathogenic role of B cells in MS might lie less with the generation of autoantibodies, and more with their capacity to interact with other immune cell types, such as T cells.

Experimental autoimmune encephalomyelitis (EAE) is an animal disease that recapitulates many of the immune aspects of MS pathogenesis. Use of this model has helped us to understand the role of T cells, and CD4^+^ T cells in particular, in the initiation and maintenance of autoreactive inflammation in the CNS (15). However, studies using the transgenic IgH^[MOG]^ mouse strain have indicated that B cells may also play a crucial role in EAE pathology (16–18). These mice (also known as “Th”) express a knocked-in IgH chain derived from a MOG-specific antibody; thus, around 30% of their B cells are therefore specific for MOG protein (16). IgH^[MOG]^ animals develop severe EAE when immunized with either whole MOG protein (16) or with its extracellular domain (MOG_[1-125]_) (19), indicating an important role for MOG-reactive B cells in neuroimmune processes. Further, they develop spontaneous EAE when crossed to myelin peptide-specific T cell receptor transgenic lines on the B6 (17) and SJL/J (18) backgrounds. Indeed, while “T cell help” is required for the full activation of B cells in the majority of cases, there is increasing evidence that B cells can reciprocally promote effector T cell responses, notably those of the Th17 lineage (20–24). However, the potential mechanisms by which MOG-reactive B cells facilitate T cell-driven pathogenicity, such as in class II-restricted peptide immunization models of EAE, remain incompletely understood.

Here, we studied the co-operative role of B cells and T cells in CNS autoimmunity using IgH^[MOG]^ mice on the non-obese diabetic (NOD) genetic background on which EAE develops with a relapsing/chronic disease pattern (25).When immunized with the T cell-obligate peptide MOG_[35-55]_, IgH^[MOG]^ mice developed a rapid, severe form of EAE characterized by meningeal clusters of CD4^+^ T cells, B cells and CD11c^+^ dendritic cells (DCs) with alterations to the underlying extracellular matrix (ECM) reminiscent of tertiary lymphoid organs (TLOs). Th17 responses were specifically upregulated in the inflamed IgH^[MOG]^ CNS and, surprisingly, CNS-infiltrating B cells produced the Th17 maintenance factor IL-23. Further, PD1^+^CXCR5^-^ “T peripheral helper-like cells”, a recently described effector T cell subset that associates with B cells in the inflamed synovium in rheumatoid arthritis (RA) (26), were presented at a high frequency in the IgH^[MOG]^ CNS parenchyma and dura mater. Transcriptomics analysis of CNS-infiltrating B and Tph cells revealed that genes related to neurodegeneration and oxidative phosphorylation were upregulated in both cell types in IgH^[MOG]^ mice, suggesting that these processes contribute to the heightened pathology observed in these animals. Notably, IL-23 blockade abrogated severe disease in IgH[MOG] mice, as well as both CNS Tph accumulation and ROS production from both B cells and Tph cells. Together, our results demonstrate that MOG-specific B cells play a crucial role in augmenting CD4^+^ T cell-driven EAE in an IL-23-dependent manner.

## Results

### IgH^[MOG]^ mice develop severe EAE upon active immunization with MOG_[35-55]_

When immunized with MOG_[35-55]_, wildtype WT NOD mice display a disease course characterized by relapse remitting disease in the early phase that transitions to a chronic worsening phase in some animals (25). NOD-EAE has thus been considered a possible model of SPMS (27, 28), though others argue against this interpretation (29). We confirmed that MOG_[35-55]_-immunized WT NOD mice develop disease characterized by relatively mild symptoms over the first ∼100 days (Supplemental Figure 1A). We next compared the development of EAE between NOD-background WT and IgH^[MOG]^ mice upon MOG_[35-55]_ immunization. Both male and female IgH^[MOG]^ mice developed extremely severe disease within 25 days (Figure 1A), with a substantial frequency of these mice attaining ethical endpoints (10/14 IgH^[MOG]^ males vs. 0/14 WT males, p=0.0002; 12/16 IgH^[MOG]^ females vs. 0/16 WT females, p<0.0001). Histopathological analyses revealed increased lymphocyte infiltration and demyelination in the spinal cords of IgH^[MOG]^ mice relative to controls (Figure 1B).

**Figure 1.**
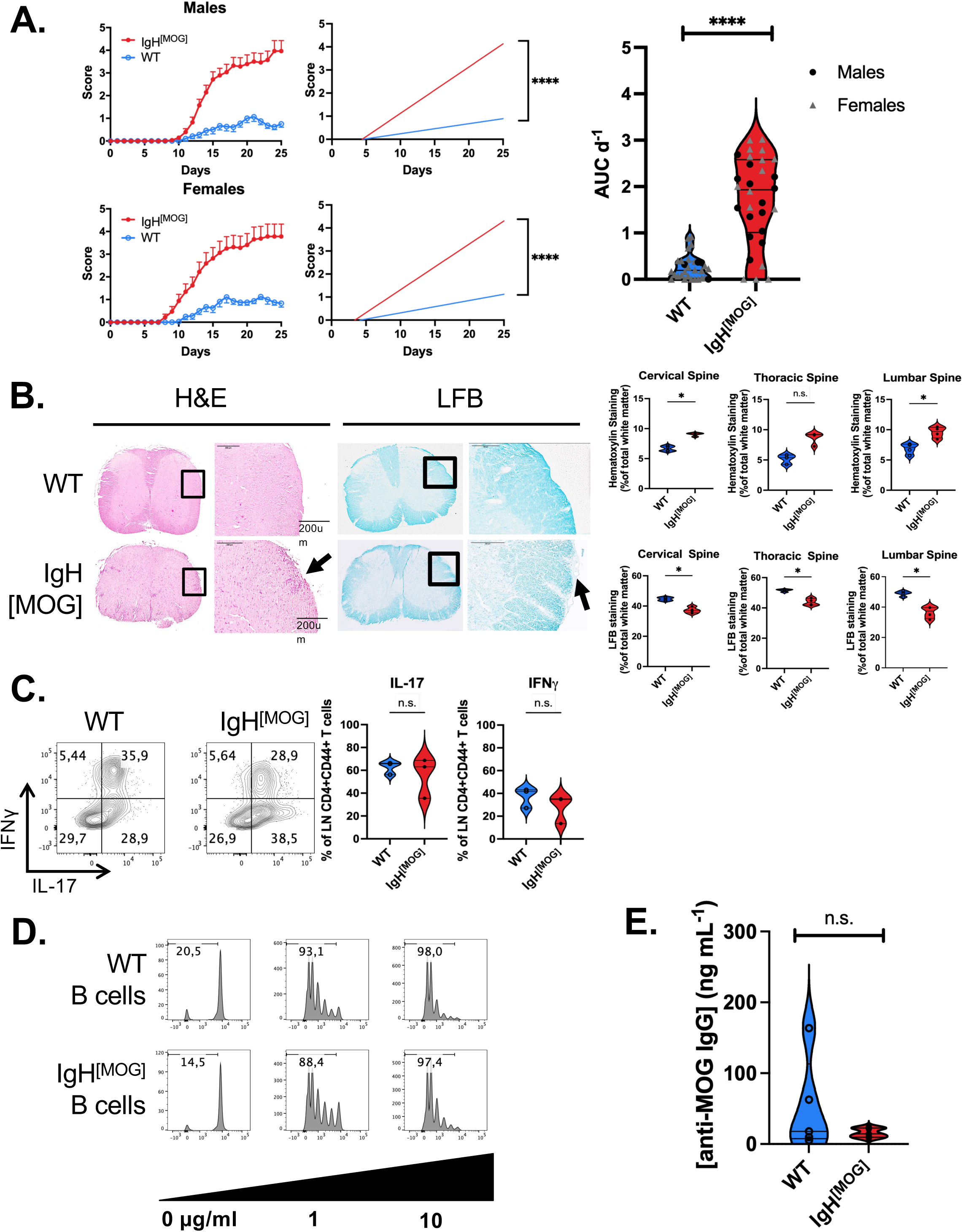
Active immunization with MOG_[35-55]_ induces severe EAE in IgH^[MOG]^ mice in the absence of increased peripheral inflammation. **A.** *Left,* Disease curves and linear regression of males (WT, n=10, IgH^[MOG]^, n=10) and females (WT, n=10, IgH^[MOG]^, n=10) immunized with MOG_[35-55]_ and monitored for development of EAE. ****, p<0.0001. Representative of 2 experiments in each sex. *Right*, AUC analysis of disease curves for all mice studied (30 WT, 30 IgH^[MOG]^). **B.** WT and IgH^[MOG]^ spinal cords were sectioned and stained with hematoxylin & eosin (H&E; immune infiltration) and Luxol fast blue (LFB; myelin). Slides were scanned at 10X magnification for quantification. WT, n=4, IgH^[MOG]^, n=4. *, p<0.05, *t*-test. **C.** WT (n=3) and IgH^[MOG]^ (n=3) mice were immunized with MOG_[35-55]_. LN cells were isolated at disease onset and IFNγ and IL-17 expression were determined by flow cytometry. Gated on live CD4^+^CD44^+^ events. **D.** WT or IgH^[MOG]^ B cells were co-cultured with CellTrace Violet-labeled MOG_[35-55]_-specific 1C6 T cells that were pulsed, or not, with the indicated concentrations of MOG_[35-55]_. Gate frequencies indicate the percentage of cells undergoing at least one cell division. Representative of triplicate cultures. **E.** Sera were collected from MOG_[35-55]_-immunized WT (n=5) and IgH^[MOG]^ (n=5) mice, and concentration of MOG-specific IgG was assessed by ELISA. n.s., not significant; *, p<0.05; ****, p<0.0001; linear regression analysis (**A**), *t*-test (**A-C, E**).

We next examined peripheral T cell responses prior to disease onset. No differences in IL-17 or IFNγ were detected by either *ex vivo* flow cytometry (Figure 1C) or by ELISA after MOG_[35-55]_ peptide recall (Supplemental Figure 1B). Next, to identify a possible role for IgH^[MOG]^ B cells in directly driving T cell responses, we pulsed IgH^[MOG]^ or WT B cells with MOG_[35-55]_ and co-cocultured them with MOG_[35-55]_-specific 1C6 transgenic T cells (30–32). However, no differences were observed in T cell proliferation (Figure 1D), indicating that IgH^[MOG]^ B cells were not intrinsically better at presenting antigen to T cells.

Furthermore, while the cervical LNs are a potential site of T cell reactivation in MS (33), we observed only limited production of IL-17 and IFNγ in this compartment. While a large proportion of CD4^+^ T cells in cervical LNs were positive for TNFα, no differences were detected between WT and IgH^[MOG]^ (Supplemental Figure 1C).

Antigen-specific antibody (Ab) secretion is the primary function of B cells. Further, oligoclonal immunoglobulin (Ig) banding in cerebrospinal fluid (CSF) is an important diagnostic marker for MS (34). However, no differences in MOG-specific circulating IgG were found in IgH^[MOG]^ serum relative to controls (Figure 1E), suggesting that severe disease in transgenic animals was not accompanied by an increase in MOG-specific autoantibodies. Together, these data showed that the presence of myelin-reactive B cells can exacerbate CNS autoimmunity and tissue damage when EAE is induced in a CD4^+^ T cell dependent manner. However, increased disease severity did not appear to be accompanied by markers of peripheral T and B cell activation.

### Accumulation of meningeal TLOs in IgH^[MOG]^ mice

Target organ-infiltrating immune cells are essential to the pathogenesis of autoimmune disease. We therefore enumerated the frequency and absolute number of CNS-infiltrating immune cells − CD19^+^ B cells, CD4^+^ T cells, CD8^+^ T cells, CD11c^+^ dendritic cells (DCs), CD11b^+^CD11c^-^ macrophages and Ly6G^+^ neutrophils − in immunized WT and IgH^[MOG]^ mice. The relative frequencies (Figure 2A) and absolute numbers (Supplemental Figure 2) of CD4^+^ T cells, CD8^+^ T cells, DCs, macrophages and neutrophils were significantly increased in the CNS of IgH^[MOG]^ mice relative to WT; however, B cell frequency and absolute numbers were unchanged.

**Figure 2.**
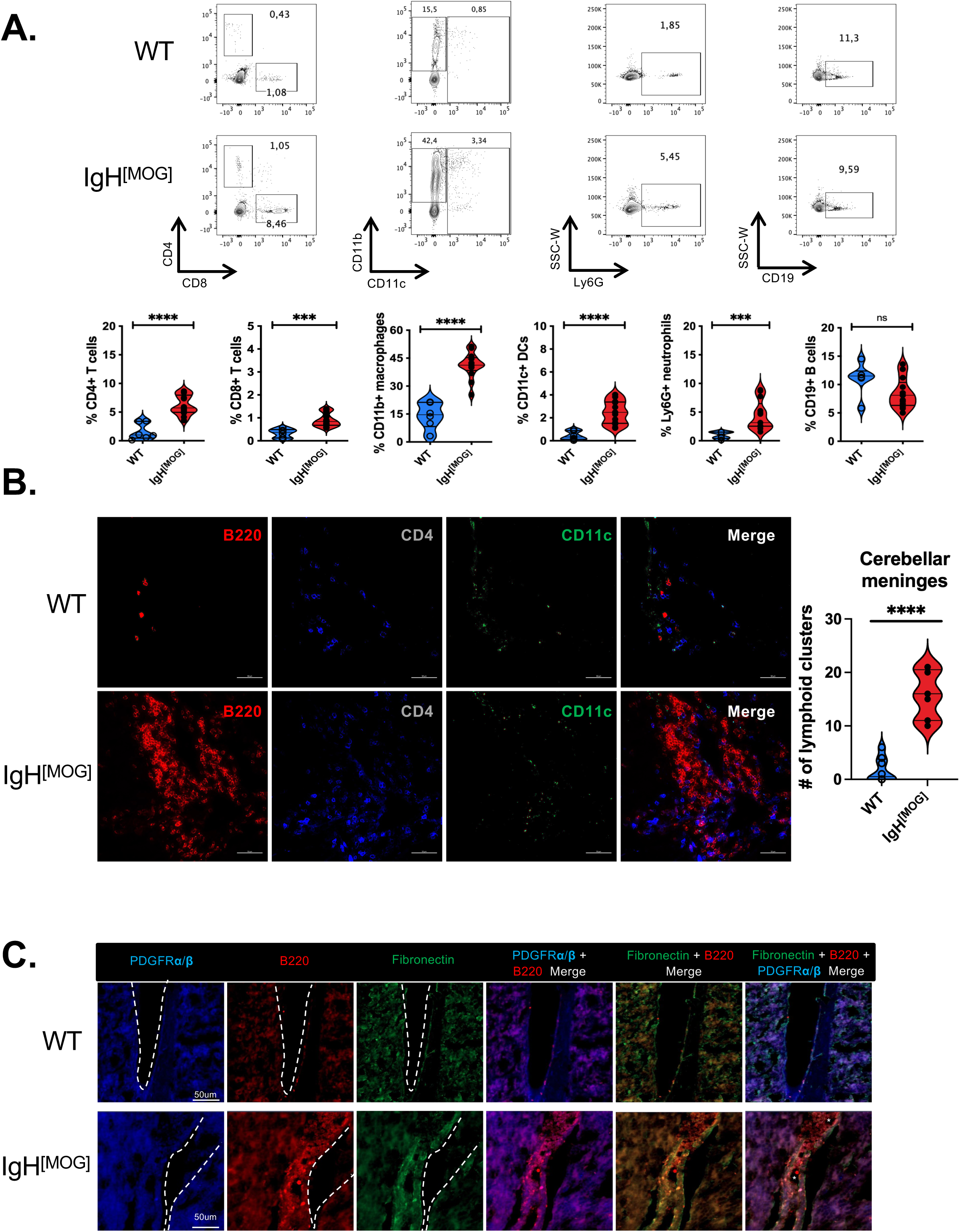
Increased lymphocytic aggregation in the meninges of immunized IgH^[MOG]^ mice. **A.** Mononuclear cells were isolated from the CNS of immunized WT (n=6) and IgH^[MOG]^ (n=12) mice. Frequency of CD19^+^ B cells, CD4^+^ and CD8^+^ T cells, Ly6G^+^ neutrophils, CD11b^+^CD11c^-^ macrophages and CD11c^+^ dendritic cells were enumerated by flow cytometry. **B.** Brain and spinal cord meningeal sections from MOG_[35-55]_-immunized WT (n=7) or IgH^[MOG]^ (n=7) mice were assessed by immunofluorescence (IF) for expression of B220 (red), CD4 (blue), and CD11c (green), and lymphocyte aggregates from the cerebellar meningeal compartment were enumerated. **C.** Assessment of the ECM near meninges in the cerebellum was determined by IF. Expansion on the meningeal ECM is shown by fibronectin (green) and PDGFRα/β (blue) which overlap with B220 B cell clusters. Dotted lines outline the meningeal membrane and (*) denotes vascular endothelium.

Meningeal TLOs have been documented in progressive MS and their presence correlates with poor outcomes (7, 35). Their presence in EAE is dependent on murine genetic background and the manner of disease induction; while TLOs arise upon active immunization of SJL/J mice with PLP_[139-151]_ (36), they do not appear in MOG_[35-55]_-immunized C57BL/6J mice, rather arising only upon adoptive transfer of Th17 cells (37). Whether or not they are a feature of NOD-EAE is as-yet unknown. As B cells are crucial to TLO formation (37), we assessed their presence in WT versus IgH^[MOG]^ mice. Strikingly, we observed clustering of CD4^+^ T cells, B cells and CD11c^+^ DCs of varying magnitude in the CNS of IgH^[MOG]^ relative to WT mice (Figure 2B).

TLOs are defined as tissue immune cell aggregates organized within a stroma cell network in inflamed non-lymphoid tissues(38). We found an increase of elaborate stroma network near the meninges in IgH^[MOG]^ mice as demonstrated by the organized expansion of fibronectin, PDGFRαβa/b IF staining underneath the B cell clustering (Figure 2C). Together, these data demonstrate that TLOs accumulate in the meninges of MOG_[35-55]_-immunized IgH^[MOG]^ mice.

### The IgH[MOG] CNS is characterized by inflammatory B cells and Th17 cells

As we did not see an increase in frequency or absolute number of B cells in the CNS of immunized IgH^[MOG]^ mice (Figure 2A), we next assessed whether there might be differences in the proportion of functional B cell subsets between the groups. Indeed, we observed a stark increase in the frequency of class-switched (CS) IgM^-^IgD^-^ B cells in the CNS (Figure 3A) of IgH^[MOG]^ mice, which was accompanied by a decrease in the frequency of immature and mature B cells. In line with our finding that serum MOG-specific IgG was comparable between WT and IgH^[MOG]^, we observed no differences in B cell subpopulation frequency in the spleen (Supplemental Figure 3A). Further, we observed no difference in MHC class II expression between CNS-infiltrating WT and IgH^[MOG]^ B cells (Supplemental Fig 3B). On the other hand, an increase in GL-7^+^Fas^+^ germinal center (GC) B cells was seen in the IgH^[MOG]^ CNS (Supplemental Figure 3C).

**Figure 3.**
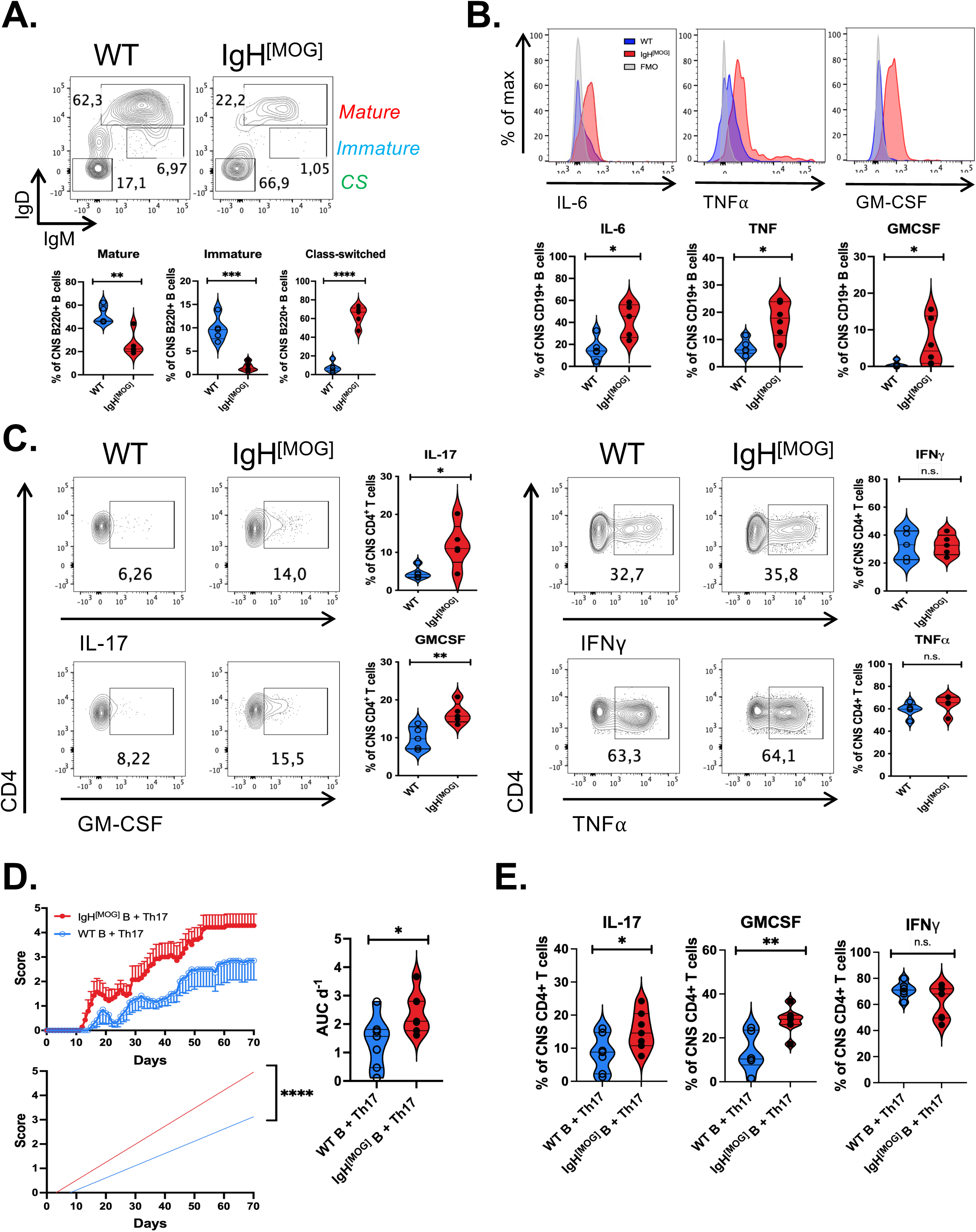
Inflammatory B cells facilitate Th17 responses in IgH^[MOG]^ mice. **A.** The frequency of immature (IgM^mid^IgD^mid^), mature (IgM^hi^IgD^hi^) and class-switched (CS; IgM^-^IgD^-^) B cells were assessed from WT (n=5) and IgH^[MOG]^ (n=5) CNS. Gated on live B220^+^ events. **B.** Expression of the indicated cytokines was assessed from WT (n=5) and IgH^[MOG]^ (n=5) CNS B cells. Gated on live CD19^+^ events. **C.** WT (n=5) and IgH^[MOG]^ (n=5) were immunized with MOG_[35-55]_ and CNS-infiltrating CD4^+^ T cells were isolated at endpoints. Expression of the indicated cytokines was assessed by intracellular flow cytometry. Gated on live CD4^+^ events. **D**. NOD.*Scid* mice were passively infused with 2×10^6^ WT or IgH^[MOG]^ B cells, and received 2×10^6^ 1C6 Th17 cells 7 days later. Mice were monitored for signs of EAE. *Left*, disease curve from one of 2 transfer experiments; *right*, AUC comparison of mice from both transfers (n=7, both groups of recipient mice). **E**. CNS-infiltrating CD4^+^ T cells were isolated from adoptive transfer recipients in (**D**) and the indicated cytokines were assessed by flow cytometry. Gated on live CD4^+^ events. n.s., not significant; *, p<0.05; **, p<0.01; ***, p<0.001; ****, p<0.0001; *t*-test (**A-E**); linear regression (**D**).

B cells can generate canonical inflammatory cytokines such as IL-6 (ref (39)), TNFα (40) and GM-CSF (41), with their production of such cytokines being linked to worsened outcomes in MS. We observed that a significantly greater proportion of IgH^[MOG]^ CNS B cells produced these cytokines as compared to WT counterparts, with the differences in IL-6 and TNFα being most apparent (Fig 3B).

Next, as we had observed an elevated frequency of CD4^+^ T cells in the CNS of sick IgH^[MOG]^ mice, we next examined the capacity of these cells to produce inflammatory Th1 and Th17 cytokines by flow cytometry, due to the well-established role of these CD4^+^ effector T cell subsets in EAE (42). We observed no differences between IgH^[MOG]^ and WT CD4^+^ T cells in their production of IFNγ and TNFα in the CNS. By contrast we saw a strong upregulation of IL-17 production from CNS-infiltrating CD4^+^ T cells from IgH^[MOG]^ at disease endpoint (Figure 3C). Notably, production of GM-CSF, a key pathogenic cytokine implicated in Th17-driven tissue inflammation, was also augmented in IgH^[MOG]^ CD4^+^ T cells.

B cells have been reported as promoting Th17 differentiation (21); further, both the congenital absence (22) and therapeutic depletion (23, 24) of B cells correlates with diminished circulating Th17 cell frequency. Thus, to test whether IgH^[MOG]^ B cells facilitate Th17 responses *in vivo*, we adapted our previously described Th17 adoptive transfer protocol (32) by seeding NOD.*Scid* mice with immunologically naïve WT or IgH^[MOG]^ B cells, and subsequently injecting MOG_[35-55]_-specific 1C6 transgenic (30) Th17 cells. Mice that had been infused with IgH^[MOG]^ B cells developed substantially more severe Th17-driven EAE than those reconstituted with WT B cells (Figure 3D). This was characterized by an increased frequency of IL-17^+^ and GM-CSF^+^ CNS-infiltrating Th17 cells in IgH^[MOG]^ B cell-reconstituted hosts (Figure 3E, Supplemental Figure 3D). Taken together, these data show that severe EAE in IgH^[MOG]^ mice is characterized by the presences of inflammatory B cells and Th17 cells, and that IgH^[MOG]^ B cells can promote Th17-mediated disease processes.

### Augmented IL-23 production by IgH^[MOG]^ mice leads to exacerbated disease

These findings led us to investigate whether IgH^[MOG]^ B cells may be characterized by increased production of the Th17 stabilization factor IL-23, which is essential to Th17 pathogenicity *in vivo* (43, 44) and which can be secreted by human B cells (45). The frequency of B cells co-expressing both chains of IL-23 (p40^+^p19^+^) was strongly upregulated in the IgH^[MOG]^ CNS, despite showing no differences in co-expression of the chains of the related cytokine, and Th1 differentiation factor, IL-12 (p40^+^p35^+^) (Supplemental Figure 4A). Further, no differences in the co-expression of p19 and p40 were observed between WT and IgH^[MOG]^ DCs, astrocytes (46) or microglia (Supplemental Figure 4B). Use of an antibody that detected IL-23 heterodimer confirmed that CNS IgH^[MOG]^ B cells show higher expression of this cytokine (Figure 4A). As we had observed an increase in B cell-rich aggregates in the IgH^[MOG]^ meninges, we assessed if these B cells were also a source of IL-23. Indeed, immunofluorescence staining showed co-localization of IL-23 with B cells in the IgH^[MOG]^ meninges (Figure 4B).

**Figure 4.**
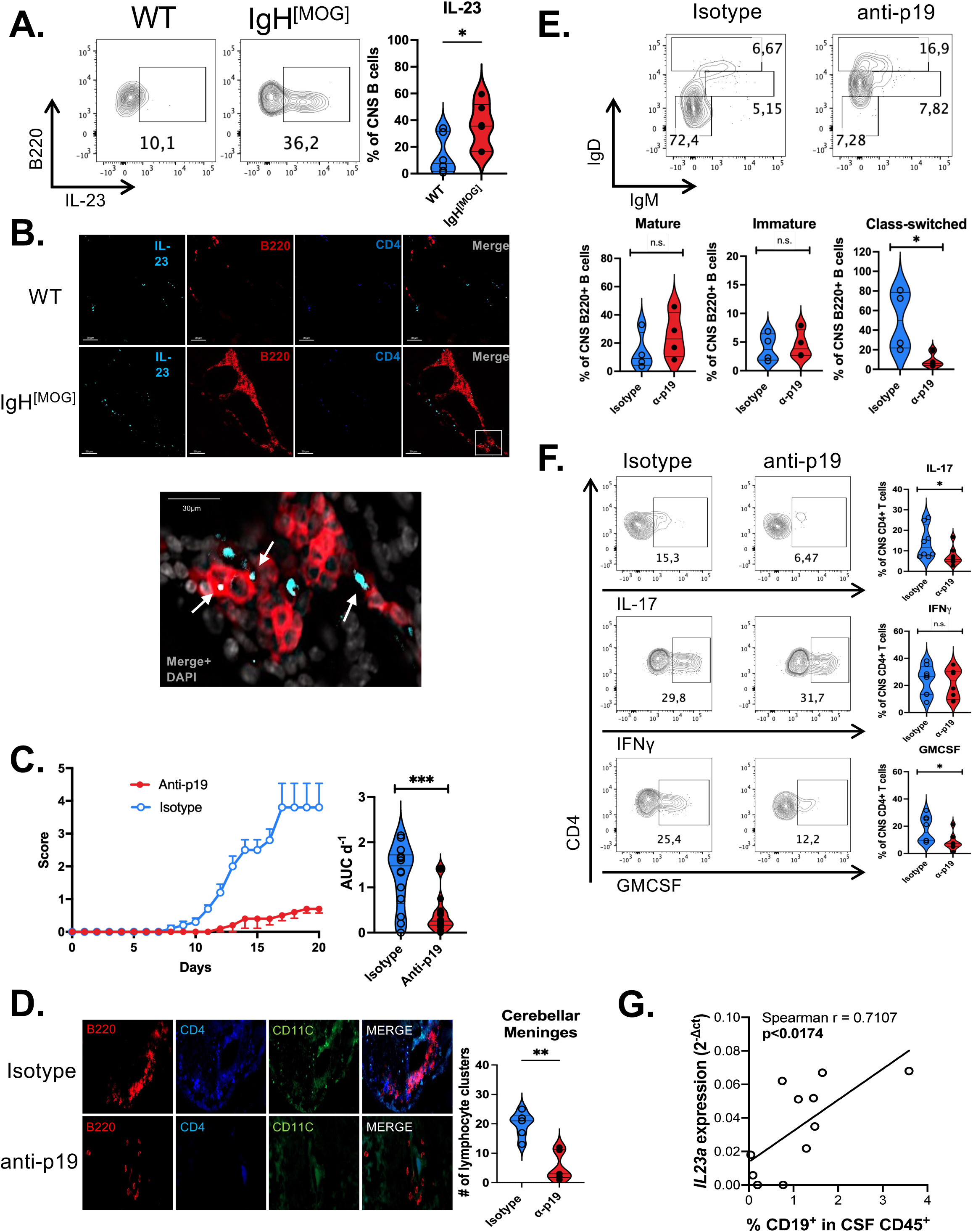
CNS IgH^[MOG]^ B cells upregulate IL-23. **A.** NOD (n=6) and IgH^[MOG]^ (n=6) mice were immunized with MOG_[35-55]_ and B cells were isolated from CNS tissues at endpoints. Expression of IL-23 heterodimer was assessed by intracellular flow cytometry. **B.** Brain and spinal cord meningeal sections from MOG_[35-55]_-immunized WT (n=3) or IgH^[MOG]^ (n=3) mice were assessed by immunofluorescence (IF) for expression of IL-23 (aqua), B220 (red), CD4 (blue). **C.** IgH^[MOG]^ mice were immunized with MOG_[35-55]_ and were treated on day (-1) and day 6 with 1 mg anti-p19 (n=5) or rIgG1 isotype (n=5). Disease curves representative of 3 experiments. AUC calculations include all mice studied; anti- p19 (n=14) or rIgG1 isotype (n=14). **D.** Cerebellar meningeal sections from MOG_[35-55]_-immunized IgH^[MOG]^ mice, treated with isotype (n=6) or anti-p19 (n=6), were assessed for lymphocyte clusters. **E.** CNS-infiltrating mature, immature and class-switched B cells were assessed by flow cytometry from anti-p19 (n=4) or rIgG1 isotype (n=4)-treated immunized IgH^[MOG]^ mice. Gated on live B220+ events. **F.** CD4^+^ T cells were isolated from p19 (n=5) or rIgG1 isotype (n=5)- treated immunized IgH^[MOG]^ mice and the indicated cytokines were assessd by flow cytometry. **G.** *Il23a* was assessed from CSF immune cells of MS-affected individuals (10 RRMS, 1 PPMS; 9 females, 2 males) by qPCR and was correlated to the frequency of CD19^+^ B cells within the CD45^+^ CSF leukocyte population of each patient. n.s., not significant; *, p<0.05; **, p<0.01; ***, p<0.001, *** *, p<0.0001; *t*-test (**A-F**); linear regression (**G**).

The robust upregulation of IL-23 by B cells in the IgH^[MOG]^ CNS led us to ask whether it might be essential for the exacerbated disease seen in these animals. We therefore actively immunized IgH^[MOG]^ mice and administered either anti-IL-23p19 blocking antibody (47) or isotype control. *In vivo* blockade abrogated severe EAE (Figure 4C), characterized by reduced lymphocyte infiltration and demyelination in the spinal cord (Supplemental Figure 4D). As Th17 responses are crucial to TLO formation (37), we next examined the effects of IL-23 blockade on the presence of TLOs in the meninges. Administration of anti-IL23p19 diminished the number of lymphocyte clusters in the cerebellar meninges of IgH^[MOG]^ mice (Figure 4D). Concomitantly, we observed a decrease in the intricate ECM network underneath, suggesting a decrease in meningeal TLOs in IgH^[MOG]^ mice after administration of anti-IL23p19 (Supplemental Figure 4C).

Turning to the impact of IL-23 in cellular inflammation in the IgH^[MOG]^ CNS, we found that the frequency of class-switched B cells was reduced in immunized animals treated with anti-p19 (Figure 4E). Further, CNS-infiltrating CD4^+^ T cells from anti-p19-treated IgH^[MOG]^ mice showed decreased IL-17 and GM-CSF expression (Figure 4F); however, no differences in IFNγ production between T cells from anti-19 or isotype-treated mice were observed.

The cerebrospinal fluid is a useful proxy for pathologic processes within the CNS parenchyma (48) and is routinely sampled in MS diagnosis. We therefore asked whether expression of IL-23p19 transcript in the CSF of people with MS (pwMS) were altered in relation to the presence of B cells. We examined the CSF of 11 pwMS for expression of *Il23a* (IL-23p19 gene name) and correlated this to the frequency of CD19^+^ B cells in the CSF of the same individuals. A robust correlation was identified (Spearman r = 0.7107) between *IL23a* and the proportion of CD45^+^ leukocytes that were CD19^+^ (Fig. 4G). In sum, these data show that excessive production of IL-23 by IgH^[MOG]^ B cells may underpin the severe pathology observed in these animal and that IL-23 transcript correlates with the presence of B cells in CNS-associated tissues from pwMS.

### T peripheral helper (Tph)-like cells accumulate in the IgH^[MOG]^ CNS

T follicular helper (Tfh) cells play a critical role in germinal center formation and B cell immune activation, and were recently shown to collaborate with B cells in driving Th17-dependent EAE (49). We therefore examined the presence of PD- 1^+^CXCR5^+^ Tfh cells in splenic and CNS tissues from WT and IgH^[MOG]^ mice. Intriguingly, while Tfh were relatively rare in all cases, we instead observed an increase in PD-1^+^CXCR5^-^ CD4^+^ T cells in the IgH^[MOG]^ CNS (Figure 5A). These cells were reminiscent of the recently discovered “T peripheral helper” (Tph) cell subset (26, 50). In contrast to the CNS, we observed no differences in the frequency of PD-1^+^CXCR5^-^ CD4^+^ T cells in the spleen (Supplemental Figure 5A). Originally discovered in the context of rheumatoid arthritis (RA), Tph are recruited to the inflamed synovium, where they associate with B cells. They are distinct from Tfh (26), and indeed we found that PD-1^+^CXCR5^-^ cells from both WT and IgH^[MOG]^ mice were negative for Bcl6 (Supplemental Figure 5B), the Tfh master transcription factor (51). While present in human RA (26, 50) and lupus (52), evidence for the accumulation of Tph in murine autoimmune disease is, thus far, limited. Interestingly the frequency of PD-1^+^CXCR5^-^ CD4^+^ T cells was higher at the peak of EAE in IgH^[MOG]^ mice as opposed to at onset or recovery (Figure 5B).

**Figure 5.**
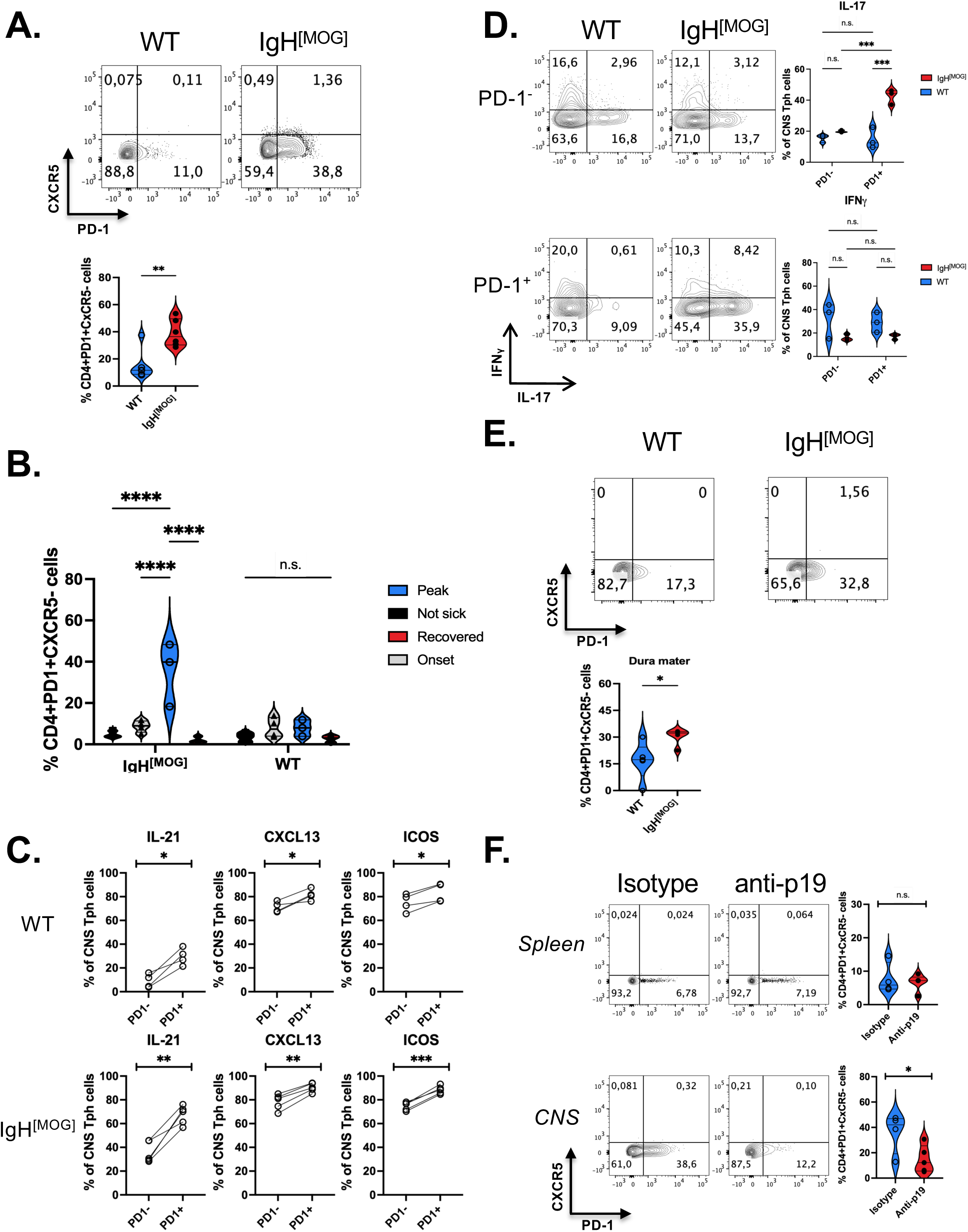
Tph cells infiltrate the IgH^[MOG]^ CNS. **A.** WT (n=6) and IgH^[MOG]^ (n=6) mice were immunized with MOG_[35-55]_ and T cells were isolated from CNS at endpoints. Expression of PD-1 and CXCR5 was assessed by flow cytometry. **B.** WT or IgH^[MOG]^ mice were immunized and sacrificed at onset (n=4 each), peak (n=3 each), recovery (n=3 each). CNS-infiltrating CD4^+^ T cells were isolated from these groups, as well as from mice that did not develop disease (n=4 each) and the frequency of PD-1^+^CXCR5^-^ cells was assessed. **C.** Expression of IL-21, CXCL13 and ICOS was assessed from CNS-infiltrating CD4^+^ T cells taken from immunized WT (n=6) and IgH^[MOG]^ (n=6) mice, and expression between PD-1^-^ and PD-1^+^ CD4^+^ subpopulations was compared. **D.** Expression of IFNγ and IL-17 was assessed from PD-1^-^ and PD-1^+^ CD4^+^ T cells from immunized WT or IgH^[MOG]^ mice. **E.** WT (n=5) and IgH^[MOG]^ (n=5) mice were immunized with MOG_[35-55]_ and T cells were isolated from the dura mater at endpoints. Expression of PD-1 and CXCR5 was assessed by flow cytometry. **F.** T cells were isolated from spleen and CNS at endpoints from anti-p19 (n=5) or isotype-treated (n=5) mice. Expression of PD-1 and CXCR5 was assessed by flow cytometry. All flow cytometry data gated on live CD4^+^ events. n.s., not significant; *, p<0.05; **, p<0.01; ***, p<0.001, ****, p<0.0001; unpaired *t*-test (**A, F**), Tukey’s multiple comparisons test (**B, D**), paired *t*-test (**C**).

In addition to being PD-1^+^CXCR5^-^Bcl6^-^, Tph are characterized as highly positive for ICOS, CXCL13 and IL-21 (26). We therefore compared expression of these markers between CNS-infiltrating PD-1^neg^ and PD-1^+^ CD4^+^ T cells of both WT and IgH^[MOG]^ mice. Expression of all three markers was upregulated in the PD-1^+^ subpopulation in both WT and IgH^[MOG]^ (Figure 5C), showing that CNS-infiltrating PD1^+^CXCR5^-^ cells express known Tph markers in the context of NOD-EAE. Interestingly, expression of IL-21 was significantly higher in IgH^[MOG]^ Tph- like cells relative to WT (Supplemental Figure 5C).

As we had found that the frequency of CNS-infiltrating IL-17^+^ CD4^+^ T cells is increased in immunized IgH^[MOG]^ mice (Figure 3C), we next asked whether production of IL-17 differed between the WT and IgH^[MOG]^ Tph populations. Indeed, CNS-infiltrating IgH^[MOG]^ CD4^+^PD-1^+^ T cells showed strikingly higher positivity for IL-17 than their WT counterparts (Figure 5D). By contrast, we saw no differences in IL-17 between WT and IgH^[MOG]^ CD4^+^PD-1^-^ T cells, nor were strain-specific differences in IFNγ seen in either the CD4^+^PD-1^+^ or CD4^+^PD-1^+^ subpopulations. Thus, Tph cells are the primary source of IL-17 from the CD4+ T cell compartment in the IgH[MOG] CNS.

Altogether, our data indicated that CD4^+^PD-1^+^CXCR5^-^ Tph cells not only accumulate in the CNS of IgH^[MOG]^ mice, but also show important phenotypic differences with WT Tph cells. To define these alterations globally, we performed mRNAseq analysis to compare the transcriptome of WT and IgH^[MOG]^ Tph cells taken from the CNS of immunized mice. We uncovered 2565 differentially expressed genes (p_adj_ < 0.05 & L_2_FC > |1.5|) of which 2154 were upregulated and 411 downregulated in IgH^[MOG]^ versus WT (Supplemental Figure 5D). In analyzing known Tph marker expression, we found that *Il21* was upregulated in IgH^[MOG]^ Tph cells relative to WT (Supplemental Figure 5E), in line with what was observed at the protein level (Supplemental Figure 5C). Further, *Cx3cr1* and *Cxcl13* were also augmented in IgH^[MOG]^ Tph cells. By contrast, *Ccr2* and *Icos* were downregulated in IgH^[MOG]^ Tph cells as compared to WT (Supplemental Figure 5E). Notably, and again mirroring our observations at the protein level (Figure 5D), expression of *Il17a*, as well as of *Il17f*, were increased in IgH^[MOG]^ Tph cells (Supplemental Figure 5F). As Tph cells were described to interact with B cells in inflammatory tissue, and having seen an increase of B cell aggregates in the meningeal TLOs, we then assessed by flow cytometry the presence of Tph cells in the dura mater. A robust upregulation of Tph cells was observed in the dura mater of IgH^[MOG]^ mice compared to WT mice (Figure 5E).

Finally, we examined whether *in vivo* depletion of IL-23 could impact Tph frequency. Indeed, the frequency of Tph was reduced in the CNS, but not spleen, of immunized IgH^[MOG]^ treated with anti-p19 (Figure 5F). Altogether, our data show that CD4^+^PD1^+^CXCR5^-^ Tph cells accumulate in the CNS of immunized IgH^[MOG]^ mice in an IL-23-dependent manner, where they are an important source of IL-17. Further, in addition to differences in their frequency, qualitative differences also exist between IgH^[MOG]^ and WT CNS-infiltrating Tph cells.

### Reactive oxygen species (ROS) generation is enhanced from IgH^[MOG]^ B cells and Tph cells

We next sought to identify molecular features, characteristic of IgH^[MOG]^ B cells, that might underpin the exacerbated disease seen in these mice. Pathway analysis of the IgH^[MOG]^ versus WT CNS-infiltrating B cell transcriptome revealed, unsurprisingly, a number of terms linked to leukocyte function and immunity. Strikingly, however, we also observed an over-representation of terms linked to neurodegeneration and oxidative stress (Figure 6A). Notably, similar terms were found to be upregulated when we compared the transcriptome of IgH^[MOG]^ CNS Tph cells to WT (Figure 6A); indeed, neurodegeneration and oxidative stress gene sets were enriched in both IgH^[MOG]^ B cells and Tph cells in the CNS (Figure 6B).

**Figure 6.**
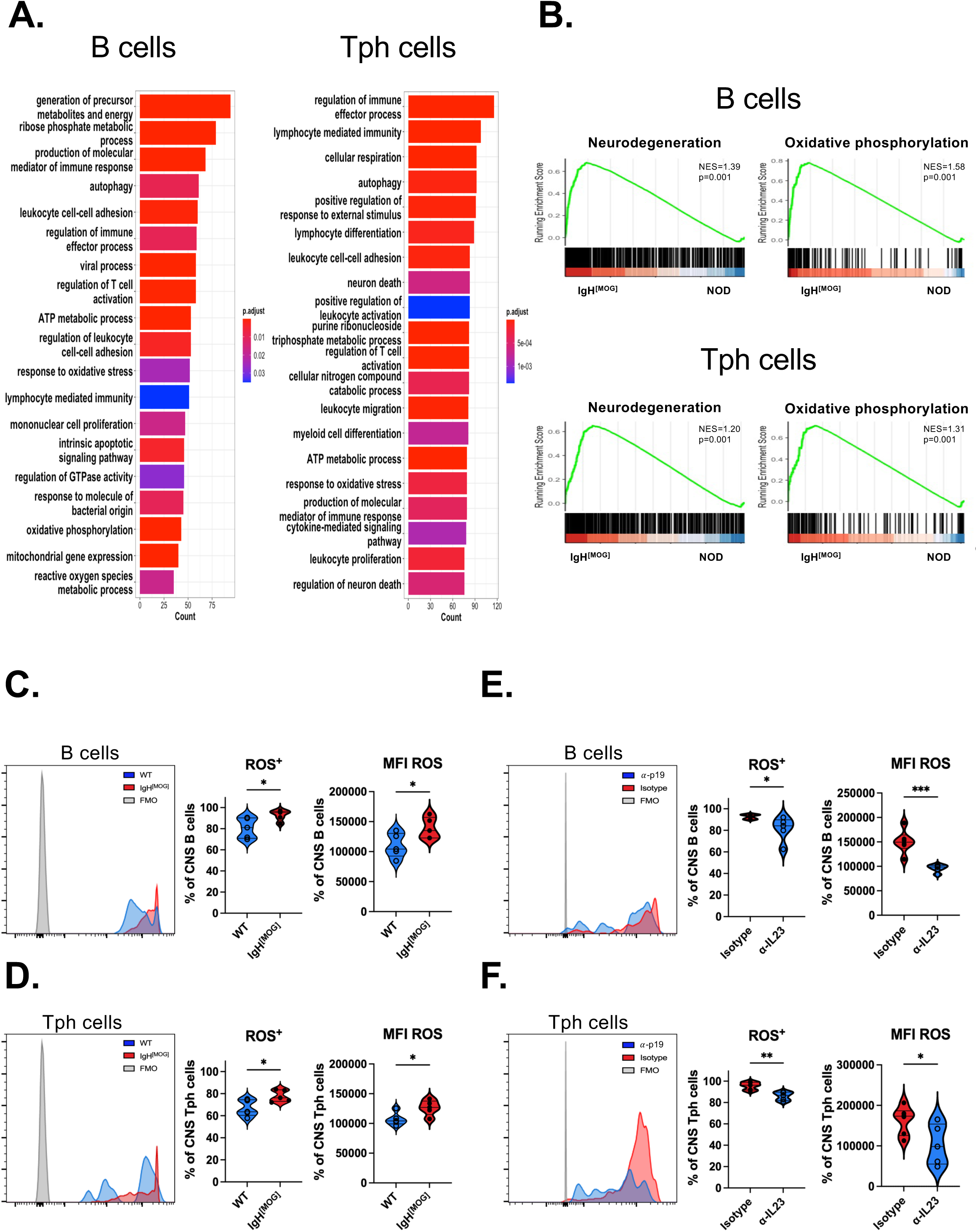
CNS-infiltrating IgH[MOG] B cells and Tph cells show increased production of reactive oxygen species. **A.** GO term analysis of the transcriptome of CNS-infiltrating B cells and Tph cells from immunized IgH^[MOG]^ (n=3) vs WT (n=3) at endpoint. **B.** Normalized enrichment scores (NES) for neurodegeneration and oxidative stress pathways from B cell and Tph cell transcriptomes. **C-F.** Uptake of ROS indicator CM-H2DCFDA from B cells (**C, E**) or Tph cells (**D, F**) from immunized IgH^[MOG]^ vs WT mice (**C, D**; n=5 each) or from immunized and anti-p19 vs isotype-treated IgH^[MOG]^ mice (E, F; n=5 each).

These findings suggested to us that an oxidative stress program might underly the EAE phenotype of IgH^[MOG]^ mice. We therefore examined whether CNS-infiltrating B cells and Tph cells could produce reactive oxygen species (ROS). Notably, IgH^[MOG]^ B cells (Figure 6C) and Tph cells (Figure 6D) had higher ROS production than their WT counterparts, as assessed CM-H2DCFDA, a fluorescent oxidative stress indicator. Then, we examined whether *in vivo* depletion of IL-23 could impact ROS production from IgH^[MOG]^ B and Tph cells. Interestingly, IL-23 blockade was decreased CM-H2DCFDA positivity from CNS-infiltrating B cells (Figure 6E) and Tph cells (Figure 6F). In sum, these data show that CNS-infiltrating IgH^[MOG]^ B cells and Tph cells mount oxidative stress responses that are IL-23 dependent.

## Discussion

The role of CD4^+^ T cells in the autoimmune pathogenesis of MS is well-established: genome-wide association studies (GWAS) have revealed that polymorphisms in the human leukocyte antigen class II region are strongly associated with MS susceptibility (53), and many of the current treatments of MS are believed to target T cell responses (54–56). However, the success of the B cell-depleting drug ocrelizumab in both RRMS and PPMS has intensified recent interest in the contribution of B cells to disease processes (57).

IgH^[MOG]^ mice were initially described on the C57BL/6 (B6) and SJL/J genetic backgrounds. On the B6 background, IgH^[MOG]^ mice showed an increased incidence of EAE, relative to WT, when immunized with whole MOG protein(16). Notably, IgH^[MOG]^ SJL/J mice developed EAE of greater severity than controls when immunized with the myelin-derived epitope proteolipid protein (PLP)_[139-154]_(16), which induces a relapsing/remitting disease pattern in SJL/J background mice(58). This suggested that the presence of MOG-reactive B cells could contribute to EAE pathology that was driven by a class II-restricted peptide. It was later shown that when IgH^[MOG]^ mice were crossed to the 2D2 MOG_[35-55]_ TcR-transgenic strain on the C57BL/6 background, the resulting double-transgenics spontaneously developed Devic’s-like disease while single transgenic mice did not (17). Similarly, IgH^[MOG]^ x TCR^1640^ (ref (18)) and IgH^[MOG]^ x 1C6 (ref (30)) double-transgenic mice, on the SJL/J and NOD backgrounds respectively, develop spontaneous EAE at a high rate of incidence(30). These findings indicated that the collaboration of both myelin-specific B and T cells in the same animal could induce CNS autoimmunity; however, it was difficult to determine whether B or T cell-driven responses were initially responsible for disease induction in this model, and thus the nature of a putative collaboration between B cells and T cells in disease processes remained incompletely defined.

In our study, we actively immunized IgH^[MOG]^ mice on NOD background with the class II-restricted peptide MOG_[35-55]_. This permitted us to study the contribution of MOG-reactive B cells in a model of EAE that is initiated by CD4^+^ T cells. While WT NOD mice developed a gradual, chronic MOG_[35-55]_-driven disease course, with advanced symptoms appearing as late as >100 days post immunization, we found that immunized IgH^[MOG]^ NOD mice develop severe disease within a matter of weeks. Thus, by using a myelin-derived, class II-restricted, immunogen, we show that B cells augment EAE even when CD4^+^ T cells initiate disease. Interestingly, no differences in circulating anti-MOG IgG were identified between immunized WT and IgH^[MOG]^, suggesting that worsened disease in IgH^[MOG]^ mice is not the result of increased production of encephalitogenic autoantibody.

The role of CD4^+^ T cells in licensing peripheral B cell responses is well-established. Th17 cells, in particular, can foster B cell responses by inducing class switching to inflammatory IgG1 and IgG2a (59) and by recruiting B cells to TLO structures (60). Here, we provide evidence that the converse is also true, and that B cells may themselves shape Th17 responses. CNS-infiltrating CD4^+^ T cells in IgH^[MOG]^ mice showed significantly higher expression of IL-17 and GM-CSF, but not of IFNγ, indicating that Th17 responses are elevated in these mice. Further, CNS-infiltrating IgH^[MOG]^ B cells showed increased expression of IL-23, an innate immune-associated cytokine that stabilizes the Th17 lineage and is required for the pathogenicity of Th17 cells (43, 44), and *in vivo* blockade of IL-23 sharply reduced EAE in IgH^[MOG]^ mice.

Overall, our findings show that B cells can play an important supportive role in facilitating pathogenic Th17 function and are consistent with a model by which bidirectional B:Th17 interactions are essential in CNS autoimmunity. Indeed, there is emerging evidence that B cells can promote Th17 responses. Both peritoneal CD5^+^B220^lo^ (ref (20)) and LPS-stimulated splenic CD80^+^CD86^+^CD44^+^ (ref (21)) B cells can elicit Th17 differentiation. Further, production of IL-17 is defective from T cells taken from agammaglobulinemia patients that lack B cells(22) and therapeutic depletion of B cells with rituximab can reduce Th17 function in the context of MS (23) as well as in rheumatoid arthritis (24). The mechanisms underpinning bidirectional B:T cell interactions remain to be fully elucidated.

Human B cells can secrete IL-23, and repression of this capacity is posited as a mechanism of action for interferon-beta in MS (45). It was previously observed that peripheral B cells from naïve IgH^[MOG]^ mice possess the capacity to secrete IL-23 (61); both upon pulsing with rMOG, and most strikingly upon co-culture with MOG_[35-55]_-specific 2D2 T cells in the presence of either rMOG or MOG_[35-55]_. B cell production of IL-23 in response to LPS is repressed by the inhibitory receptor Tim-1 (ref (62)). We now show that immunization with MOG_[35-55]_ elicits IL-23 expression from CNS-infiltrating B cells *in vivo*, and that IL-23 is essential to both EAE and TLO formation from these animals.

Using mass cytometry, Brenner and colleagues found that PD1^+^CXCR5^-^ CD4^+^ T cells accumulate in the inflamed synovium of RA patients. These Tph cells secrete B cell attractant factors such as CXCL13 and IL-21 and associate with B cells in the inflamed synovium, both in defined lymphoid aggregates as well as more diffusely. However, in contrast to Tfh cells, Tph are not defined by expression of the transcription factor Bcl6 (26). While these cells are also observed in patients with lupus (52) and ANCA-associated vasculitis (63), their potential roles in murine models of autoimmunity remain obscure. We have found an enrichment of PD1^+^CXCR5^-^ CD4^+^ T cells in the CNS, but not immune periphery, of immunized IgH^[MOG]^ mice. These cells are Bcl6^neg^ and show increased expression of the Tph markers ICOS, CXCL13 and IL-21, relative to PD-1^-^ counterparts. Intriguingly, we find that the presence of PD-1^+^CXCR5^-^ cells is highest in mice of peak EAE severity; Tph cells were previously shown to correlate with the severity of rheumatoid arthritis (26) and lupus (52). Here, we also show that Tph cells can be found in the dura mater of immunized mice and their frequency was higher in IgH^[MOG]^ mice. Further, the accumulation of Tph cells in the CNS is dependent on IL-23, revealing a heretofore unknown function of this cytokine.

Oxidative stress has been described as an important factor for chronic inflammation and different studies have identified a role for oxidative stress in MS (64, 65). We now show that Tph cells as well as B cells have a general oxidative stress programming that is IL-23 dependent.

In conclusion, we provide evidence that in the presence of a B cell repertoire that is skewed towards MOG, NOD-background mice develop unusually rapid and severe CD4^+^ T cell-mediated EAE. Disease is dependent on IL-23, which is produced by B cells and is required for the augmented Th17 responses seen in these mice. Furthermore, these mice show accumulation of TLOs in their meninges that are again dependent on the presence of IL-23. Finally, Tph cells aggregate in the CNS and the dura mater of these mice. Together, our findings support a critical role for myelin-reactive B cells in bolstering T cell-driven CNS autoimmunity in an IL-23-dependent manner.

## Methods

### Ethics

All mouse experiments and breedings were approved by the Animal Protection Committee of Université Laval (protocols 2021-820 and 2021-830, to M.R). Protocols and experiments involving human participants were approved by the Newfoundland Health Research Ethics Board (to C.S.M).

### Animals

IgH^[MOG]^ mice on the NOD background (30), and 1C6 mice (30–32), were obtained from Dr. Vijay Kuchroo (Brigham & Women’s Hospital, Boston), and were maintained at the animal facility of the Centre de recherche du CHU de Québec-Université Laval. NOD/ShiLtJ and NOD.*Scid* mice were purchased from Jackson Laboratories.

### EAE induction and scoring

WT NOD and IgH^[MOG]^ mice were immunized subcutaneously with 200µg MOG_[35-55]_ (Feldan), emulsified in incomplete Freund’s adjuvant (BD Difco) that was supplemented with 500 µg *M. tuberculosis* extract (BD Difco). On day 0 and day 2 post-immunization, mice received 200 ng pertussis toxin (List Biological Laboratories) intraperitoneally. Mice were monitored daily for signs of EAE, which were assessed using a semi-quantitative 0-5 scale: 0; no disease, 0.5; ruffled fur, 1; limp tail, 1.5; mild impairment in gait, 2; severe impairment in gait, 2.5; partial hind limb paralysis, 3; hind limb paralysis, 4; forelimb paralysis, 5; moribund (66). Pre-onset analyses were conducted a minimum of 5 days post-immunization but before the onset of symptoms. For comparison of disease burden, area under curve was calculated for individual disease curves and was divided by the number of days in the protocol (AUC d^-1^). For p19 blockade, mice were administered anti-p19 (clone G23-8) or istotype (rIgG1, clone HRPM; both BioXcell), 1 mg on d^-1^ and d^+6^ (47).

### Histopathology

Mice were euthanized and intracardially perfused with PBS. The spinal cord was extracted and fixed in 10% formalin and 30% sucrose prior to being embedded in paraffin. Five-to-seven-micron coronal sections of the spinal cords were collected on superfrost microscope slides (using microtome, Leica Biosystems HistoCore Autocut) and dried overnight on a drying bench at 37°C. Slides were stored at room temperature in a slide storage box until staining was performed. Sections for Hematoxylin and Eosin (H&E) were deparaffinated with xylene, while toluene was used for Luxol Fast Blue (LFB) sections. Histology was performed as previously described (67), using standard H&E to visualize lymphocyte infiltration and LFB to stain myelin in order to visualize areas of demyelination. Briefly, stained slides were scanned at 20x using a brightfield microscope scanner (Aperio AT2 DX System), and representative RGB images of the cervical spine were acquired. Quantification of the staining was performed on Image J (software, v 1.53K) in a blinded manner. Images were split into separate single colour channels using the Colour Deconvolution plugin. Thresholding was performed on the single colour channels for hematoxylin and LFB, and an area fraction measurement was performed on the white matter of the spinal cord. Staining is expressed as the percent of total white matter that is stained.

### Immunofluorescence

Immunofluorescence (IF) analysis was performed on flash-frozen brain tissue embedded in optimum cutting temperature medium (OCT). The tissue was cryo- sectioned (7-10um) and mounted onto positively charged slides. IF staining were performed with antibodies against surface immune antigens such as CD4, B220, CD45 and CD11c and in some cases against ECM antigens: Fibronectin, PDGFRα/β simultaneously. Slides were imaged using a widefield inverted microscope (AxioObserver 7) and analysis was done using Image J software.

### ELISA

Splenocytes and draining lymph nodes were isolated from NOD and IgH^[MOG]^ mice 5d post-immunization. Cells were cultured at a concentration of 10 × 10^6^ cells mL^-1^ in T cell media and stimulated, or not, with 10 mg mL^-1^ MOG_[35-55]_. Supernatants were collected at d5 of the culture for the analysis of T cell cytokine production using a combination of commercially available antibodies (Biolegend). Briefly, the capture antibodies used were: purified anti-mouse IFNg (clone R4-6A2) and purified anti-mouse IL-17 (clone TC11-18H10.1). The detection antibodies used were: biotin anti-mouse IFNg; clone XMG1.2, biotin anti-mouse IL-17; clone TC11-8H4. Following the incubation with the avidin-horseradish peroxidase (Biolegend) and TMB substrate (Mendel Scientific), colorimetric readings were performed using a SpectraMax i3 Microplate Reader.

### Measurement of serum immunoglobulin

Blood was collected from Igh[MOG] mice and WT mice at endpoint. Serum was collected by centrifuging blood samples at 2000g for 10min. Total anti-MOG IgG was quantified by using SensoLyte Anti-Mouse MOG(1–125) IgG Quantitative ELISA Kit (Anaspec).

### Isolation of CNS-infiltrating mononuclear cells

Mice were euthanized and perfused intracardially with PBS. Brain and spinal cord were dissected from the skull and vertebral column respectively and were prepared as previously described (66). Briefly, CNS tissues were digested with liberase (Roche) and DNAse I (Sigma) and cells were enriched using a 35% Percoll (GE Healthcare) gradient.

### Isolation of dura mater cells

Mice were injected intravenously with fluorochrome-labeled anti-CD45 antibody 15 minutes prior to euthanasia. Mice were then perfused intracardially with PBS and skull was gently removed mechanically to peel the dura from the edges of the skull cap. Samples were then incubated for 30 minutes at 37°C in digestion buffer (1mg/mL collagenase D (Sigma) and 50ug/mL of DNAse 1 (Sigma) in RPMI). Samples were mechanically dissociated through a 100um cell strainer and flowthrough was centrifuged at 300g for 5 minutes at 4°C. Finally, the supernatant was discarded, and cell pellet was resuspended for flow cytometry analysis.

### Flow cytometry

Single cell suspensions were obtained from spleens, lymph nodes and CNS of EAE mice. For detection of surface antigens, cells were stained with Fixable Viability Dye (eBioscience) and incubated with Fc Block (Biolegend) prior to staining with antibodies against surface antigens (CD45, CD4, CD8, CD19, CD11b, CD11c, B220, Ly6G, IgD, IgM, FAS, GL7, PD-1, CXCR5, ICOS, I-Ag^7^, GFAP; details in following section). For detection of intracellular cytokines, cells were first stimulated with 50ng ml^-1^ PMA (Sigma), 1µM ionomycin (Sigma) and 1µL mL^-1^ GolgiStop (BD) for 4 hours at 37°C, prior to being labeled with viability indicator, Fc Block and relevant surface antigens as above. They were then fixed and permeabilized (Fixation Buffer and Intracellular Staining Perm Wash Buffer, both Biolegend) and stained for intracellular markers (IFN-γ, IL-17A, TNF-α, IL-6, GM-CSF, IL-12p40, IL-12p35, IL-23p19, IL-23; Bcl6, IL-21, CXCL13; details in following section). CM-H2DCFDA, a general oxidative stress indicator, was used to detect total ROS production. Cells were incubated with 1 μM CM-H2DCFDA (Thermo Fisher Scientific, C6827) at 37°C for 30 minutes, followed by surface proteins staining as described above. Samples were analyzed on a FACS Aria (BD) and data were analyzed using FlowJo software (Treestar).

### Flow cytometry antibodies

The following monoclonal antibodies against mouse antigens were used: CD45, clone A20 (Biolegend); CD11b, clone M1/70 (eBioscience); CD11c, clone N418 (Biolegend); Ly6G, clone 1A8 (BD Biosciences); CD4, clone RM4-5 (eBioscience); CD8, clone 53-6.7 (Biolegend); CD19, clone 1D3 (eBioscience); B220, clone RA3-6B2 (eBioscience); IgD, clone 11-26c (eBioscience); IgM, clone 11/41 (eBioscience); FAS, clone 15A7 (eBioscience); GL7, clone GL-7 (eBioscience); PD-1, clone J43 (eBioscience), CXCR5, clone SPRCL5 (eBioscience); ICOS, clone C398.4A (eBioscience); GFAP, clone 1B4 (BD Biosciences); I-Ag^7^, clone 39-10-8 (Biolegend); IFN-γ, clone XMG1.2 (eBioscience); TNF-α, clone MP6-XT22 (eBioscience); IL-17a, clone TC11- 18H10.1 (Biolegend); GM-CSF, clone MP1-22E9 (eBioscience); IL-6, clone MP5- 20F3 (eBioscience); IL-21, clone mhalx21 (eBioscience); IL-23 heterodimer, clone IC18871P (R&D Systems); IL-23p19, clone fc23pg (eBioscience); IL- 12p35, clone 4D10p35 (eBioscience); IL-12p40, clone C17.8 (eBioscience); BCL6, clone IG191E/A8 (eBioscience); CXCL13, clone D58CX13 (eBioscience); CM-H2DCFDA, clone C6827 (Thermo Fisher Scientific).

### Immunofluorescence antibodies

The following monoclonal antibodies against mouse antigens were used: CD4 (RM4-5), B220 (RA3-6B2), CD11c (N418), CD45.2 (104), PDGFRα (APA5), and PDGFRβ (APB5), all of which are from eBioscience; and fibronectin (GW20021F, Sigma); smooth muscle actin (1A4, Sigma); IL-23 heterodimer (IC18871P, R&D Systems).

### Antigen presentation assay

Single cell suspensions were obtained from the spleens of unimmunized WT and IgH[MOG] mice. Cells were labeled with CD43 (Ly-48) Microbeads (Miltenyi), and CD43^+^ leukocytes (all leukocytes except resting B cells) were depleted on a magnetic MACS column (Miltyeni). Unlabelled CD43^-^ cells were collected and were subsequently stained with anti-mouse CD19. CD19^+^ B cells were purified using high-speed cell sorting. In parallel, CD4^+^ T cells were purified from the spleens of 1C6 mice using mouse CD4 MicroBeads (Miltenyi) and labeled with CellTrace Violet (CTV; Thermo Fisher Scientific). CTV-labeled 1C6 CD4^+^ T cells were cultured with B cells at a ratio of 1 CD4: 1 B, with 0, 1 or 10µg ml^-1^ MOG_[35-55]_ for 72 hours. CTV dilution was assessed by flow cytometry.

### Tfh culture

The protocol of Gao et al was adapted (68). Briefly, 5x10^5^ splenocytes from NOD mice were stimulated 24h with LPS (1 μg mL^-1^). 1x10^4^ MOG_[35-55]_-specific 1C6 CD4^+^CD62L^hi^ T cells were then plated on top of the splenocytes with 1μg mL^-1^ MOG_[35-55]_, plus 50 ng ml^-1^ IL-21 and 100 ng mL^-1^ of IL-6 (Tfh) or no additional cytokines (Th0) for 72h.

### Analysis of CSF from MS-affected individuals

5mL of CSF was collected from consenting adult participants with MS at The Health Sciences Centre neurology clinic in St. John’s, NL between July 2021 and May 2022. For CSF immune phenotyping, 2.5mL of CSF was centrifuged for 10 minutes at 300g and the CSF was removed. The CSF cell pellet was resuspended in 100uL of flow buffer (1% bovine albumin serum, 2mM EDTA, 2mM sodium azide in PBS). The CSF cell suspension was added to a DURAclone IM Phenotyping BASIC tube (Beckman Coulter), mixed and incubated at 4°C for 30 minutes. The cells were washed with 4mL flow buffer, centrifuged at 300g for 5 minutes, decanted and resuspended in 100μL 2% paraformaldehyde. Data was acquired from the whole sample using a Cytoflex flow cytometer (Beckman Coulter) and analysed using FlowJo software. The CD19^+^ cell count was expressed as a percentage of the total number of CD45^+^ cells.

For measurement of CSF cell *Il23a* expression, 2.5 mL of human CSF was centrifuged at 300g for 10 minutes and the CSF supernatant was carefully removed. The CSF cell pellet was resuspended in 500ul of Trizol and stored at - 80°C. Total RNA was extracted using chloroform extraction followed by an RNeasy Micro kit (Qiagen). 200ng of RNA was used to synthesize cDNA with the M-MLV Reverse Transcriptase kit (Invitrogen). *Il23a* transcript expression was quantified using *Il23a* (FAM labelled) and GAPDH (VIC labelled) Taqman probe/primer assays (Invitrogen), Fast Advanced Master mix (Invitrogen) and a ViiA7 Real-time PCR system (Applied Biosystems).

### mRNA-seq

At experimental endpoints, CNS tissues from WT and IgH^[MOG]^ mice were necropsied and mononuclear cells were isolated. Live CD19^+^ B cells and CD4^+^PD1^+^CXCR5^-^ Tph cells were purified by high-speed sorting (FACSAria II) and were directly processed using NEBNext Single Cell/Low Input RNA Library Prep Kit (New England Biolabs), according to the manufacturer’s instructions, for mRNA sequencing library preparation. Total mRNA was enriched using poly(A)+ primers and template-switching oligo synthesize double-stranded cDNA with reverse transcriptase. A PCR amplification of 21 cycles was performed to increase cDNA yields followed by a purification step with AxyPrep Mag PCR Clean-up kit (Axygen, Big Flats, NY, USA). Enzymatic fragmentation of cDNA and end-repair steps were followed by ligation of NEBNext adaptors and PCR enrichment step of 11 cycles to incorporate specific indexed adapters for the multiplexing. The quality of final amplified libraries was examined with a DNA screentape D1000 on a TapeStation 2200 and the quantification was done on the QuBit 3.0 fluorometer (ThermoFisher Scientific, Canada). Subsequently, mRNA- seq libraries with unique dual indexes were pooled together in equimolar ratio and sequenced for paired-end 100 pb sequencing on NovaSeq 6000 at the Next- Generation Sequencing Platform, Genomics Center, CHU de Québec-Université Laval Research Center. The average insert size for the libraries was 253 bp. The mean coverage/sample was 24M paired-end reads.

### Bioinformatics analyses

Paired-end mRNAseq fastq datasets were trimmed with trimmomatic v0.39 using the following command: *java -jar trimmomatic-0.39.jar PE -phred 33 sample_R1.fastq sample_R2.fastq trimmed/paired_sample_R1.fastq trimmed/unpaired_sample_R1.fastq trimmed/paired_sample_R2.fastq trimmed/unpaired_sample_R2.fastq ILLUMINACLIP:TruSeq3-PE.fa:2:30:10 LEADING:30 TRAILING:30 SLIDINGWINDOW:4:15 MINLEN:30* Trimmed fastq files were mapped on the *M. musculus* genome (Ensemble; Mus_musculus.GRCm39.cds.all.fa.idx) and quantified using kallisto 0.44.0. Differential expression between WT and IgH^[MOG]^ conditions cells was performed using DESeq2 v1.38.3. For identifying relevant pathways, we used Gene Ontology enrichment analysis. We focused on biological processes on all significantly differentially expressed genes using Benjamini-Hochberg method to adjust for the false discovery rate. We also used the simplify function from ClusterProfiler package to remove redundant terms. Finally, GSEA from the Broad Institute of UCSD was used to calculate the enrichment score of different gene sets.

### Statistical analyses

Comparisons between two groups were made by *t*-test. Fisher’s exact test was used to test for differences in the frequency of mice attaining ethical endpoints. For calculation of Tph frequency at different disease stages, Tukey’s multiple comparisons test was used. Two-tailed analyses were used in all instances. All error bars represent s.e.m. All statistical analyses were conducted using Prism software (GraphPad).

## Abbreviations

Ab: antibody
CNS: central nervous system
CSF: cerebrospinal fluid
CTV: CellTrace Violet
DC: Dendritic cell
DEG: differentially expressed gene
EAE: experimental autoimmune encephalomyelitis
GC: Germinal center
GM-CSF: granulocyte and macrophage colony stimulating factor
H&E: hematoxylin & eosin
IFN: interferon
Ig: immunoglobulin
IL: interleukin
LFB: Luxol fast blue
MHC: major histocompatibility complex
MOG: myelin oligodendrocyte glycoprotein
MS: multiple sclerosis
NOD: nonobese diabetic
PLP: proteolipid protein
preonset: PO
RR: relapsing/remitting
SP: secondary progressive
Tfh: T follicular helper
TLO: Tertiary lymphoid organ
TNF: tumor necrosis factor
Tph: T peripheral helper
WT: wildtype.

## Declarations

### Authors’ contributions

MRF directed the project conducted experiments and helped write the manuscript. PMIAD, NF, IA, APY and JB conducted experiments. RP, AR, BM, SL and OR conducted histological analyses. PG and NB assisted with serum detection of immunoglobulins. CJ-B and AD assisted with bioinformatics analyses. CSM contributed to writing of the manuscript. MR supervised the project and wrote the manuscript. All authors read and approved the final manuscript.

## Acknowledgements

We thank Vijay Kuchroo for providing us with NOD-background IgH^[MOG]^ and 1C6 mice respectively. We thank Françoise Morin for critical discussions; Vincent Desrosiers, Alexandre Brunet, Stéphanie Fiola for technical assistance with flow cytometry; and Kim Larose, Andrée Brisson, Mathieu Vallière-St-Amant, Cindy Ouellet and the veterinary service of Université Laval for technical assistance and collaboration regarding animal care.

## Data availability statement

All datasets will be made upon reasonable request to the corresponding author.

---

**Supplemental Figure 1. A.** WT NOD mice (n=5) were immunized with MOG_[35-55]_ and were monitored for signs of EAE for 120d. **B.** Splenic and LN cells were isolated from immunized WT (n=3) and IgH^[MOG]^ (n=3) cells at disease onset, and were stimulated with 10 μg mL^-1^ MOG_[35-55]_. Supernatants were collected at d5 and IFNγ and IL-17 were assessed by ELISA. **C.** WT (n=8) and IgH^[MOG]^ (n=8) mice were immunized with MOG_[35-55]_, and CD4^+^ T cells were isolated from cervical LNs at endpoints and assessed for expression of the indicated cytokines.

**Supplemental Figure 2.** Mononuclear cells were isolated from the CNS of immunized WT (n=6) and IgH^[MOG]^ (n=12) mice. Frequency and absolute numbers of CD19^+^ B cells, CD4^+^ and CD8^+^ T cells, Ly6G^+^ neutrophils, CD11b^+^CD11c^-^ macrophages and CD11c^+^ dendritic cells were enumerated by flow cytometry.

**Supplemental Figure 3. A.** Splenic immature, mature and CS B cells were assessed from the same mice as in **Figure 3A**. **B.** MHC class II (I-Ag^7^) expression was assessed from CNS B cells at endpoint from MOG_[35-55]_- immunized NOD (n=5) or IgH^[MOG]^ (n=5) mice. **C.** Germinal center B cells (GL- 7^+^Fas^+^) were assessed from the spleens and CNS of MOG_[35-55]_-immunized NOD (n=5) or IgH^[MOG]^ (n=5) mice. **D.** Representative plots for the data presented in **Figure 3E**. *, p<0.05; ***, p<0.001, n.s., not significant.

**Supplemental Figure 4. A.** WT (n=10) and IgH^[MOG]^ (n=10) mice were immunized with MOG_[35-55]_ and B cells were isolated from CNS tissues at endpoints. Expression of IL-12p35, IL-12p40 and IL-23p19 were assessed by intracellular flow cytometry. **B.** Dendritic cells (CD11c^+^), astrocytes (GFAP^+^), resting microglia (CD45^dim^CD11b^dim^) and activated microglia (CD45^hi^CD11b^hi^) were assessed for expression of p19 and p40 from the CNS of MOG_[35-55]_- immunized WT (n=5) or IgH^[MOG]^ (n=5) mice. DCs, microglia; n=5 each group; astrocytes, n=10 each group. **C.** ECM deposition underneath meningeal lymphoid aggregates was assessed from the samples in **Figure 4D**. **D**. Inflammatory foci (H&E) and demyelination (LFB) were quantified from cervical spine of immunized IgH^[MOG]^ mice treated with anti-p19 (n=3) or rIgG1 isotype (n=3).

**Supplemental Figure 5. A.** Splenic CD4^+^ T cells were assessed for expression of PD-1 and CXCR5 from the same mice as in **Figure 5A**. **B.** Expression of Bcl6 was assessed from WT or IgH^[MOG]^ PD-1^+^CXCR5^-^ CNS-infiltrating T cells versus *in vitro* generated 1C6 Tfh or Th0. **C.** Expression of IL-21 was compared between WT (n=5) and IgH^[MOG]^ (n=5) CNS-infiltrating CD4^+^ T cells. All flow cytometry data gated on live CD4^+^ events. n.s., not significant; ***, p<0.001; *t*-test. **D.** Volcano plot of mRNA-seq data depicting DEG (1.5x-fold increase, p_adj_<0.05) between WT (n=3) and IgH^[MOG]^ (n=3) CNS-infiltrating PD1^+^CXCR5^-^CD4^+^ T cells. The top 10 DEG are labeled on the plot. **E.** Normalized counts for Tph markers *Il21*, *Cx3cr1*, *Cxcl13*, *Ccr2*, *Icos* between WT and IgH^[MOG]^ Tph from mRNAseq. **F.** Normalized counts for Il17a, Il17f. p-values from mRNA-seq data are p_adj_.

