## Supplemental Figs 1-5 for "Myelin-reactive B cells exacerbate the severity of CD4^+^ T cell-driven CNS autoimmunity in an IL-23-dependent manner"

### Slide 1
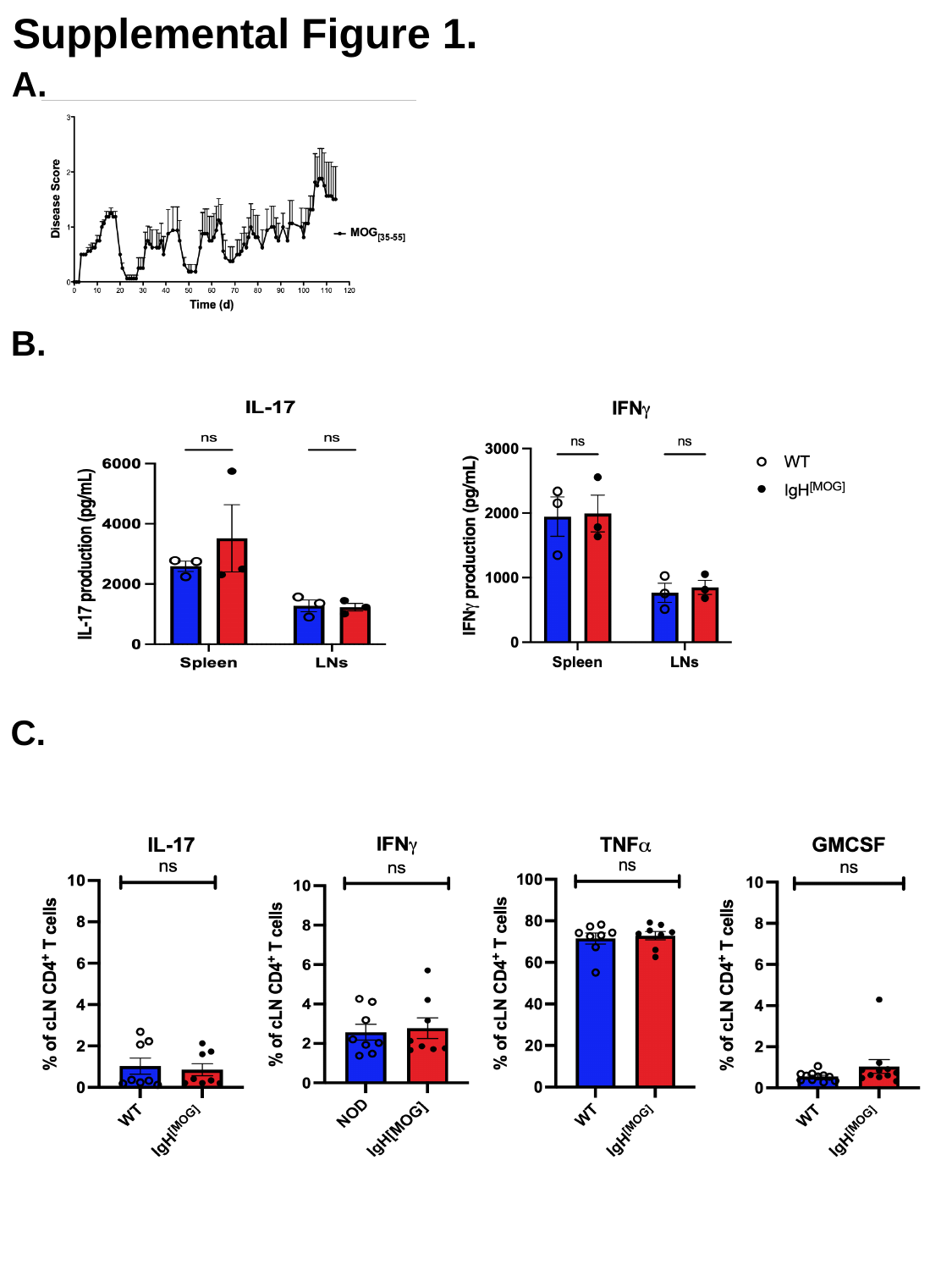

Supplemental Figure 1.
A.
B.
C.

### Slide 2
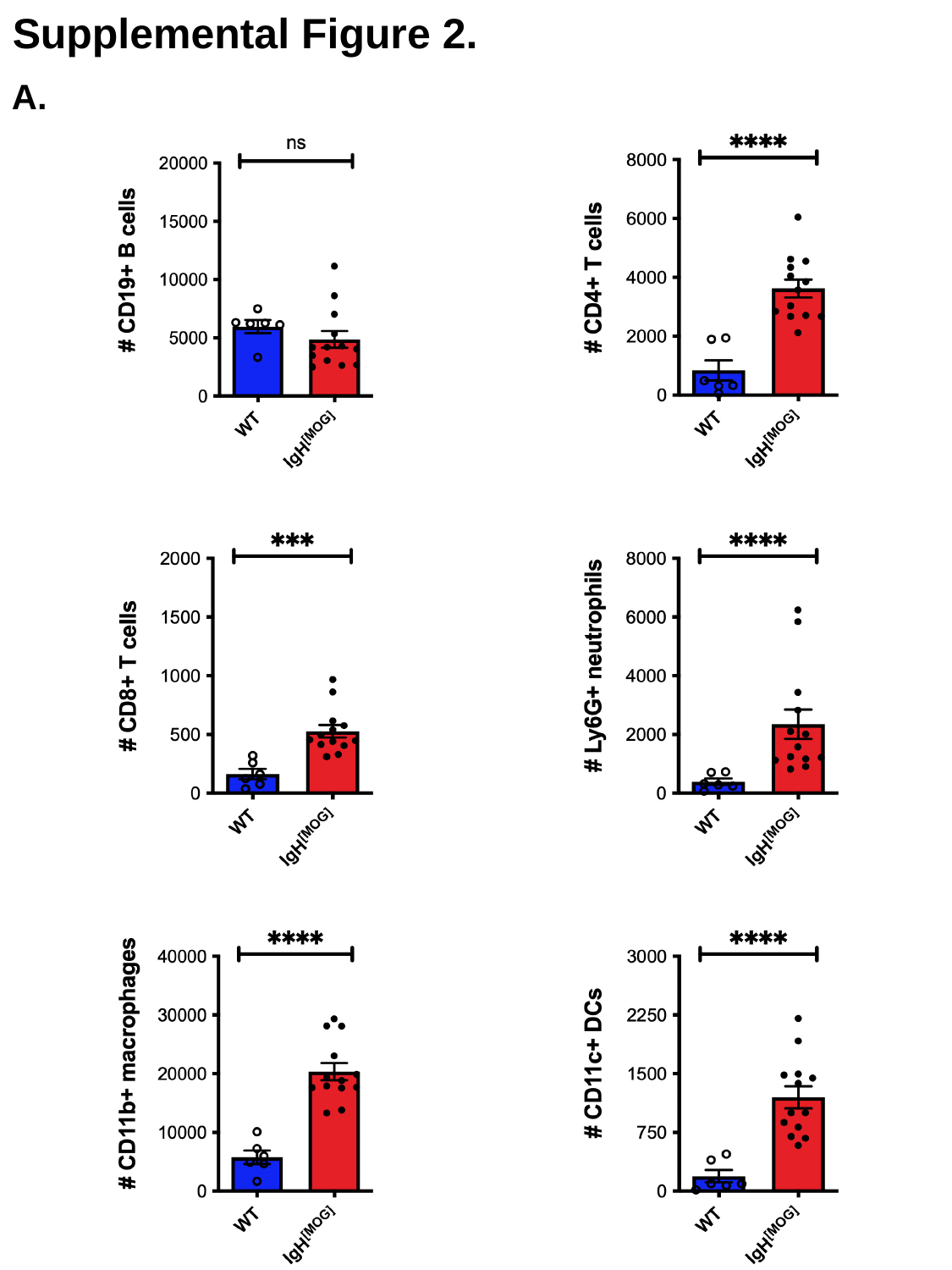

Supplemental Figure 2.
A.

### Slide 3
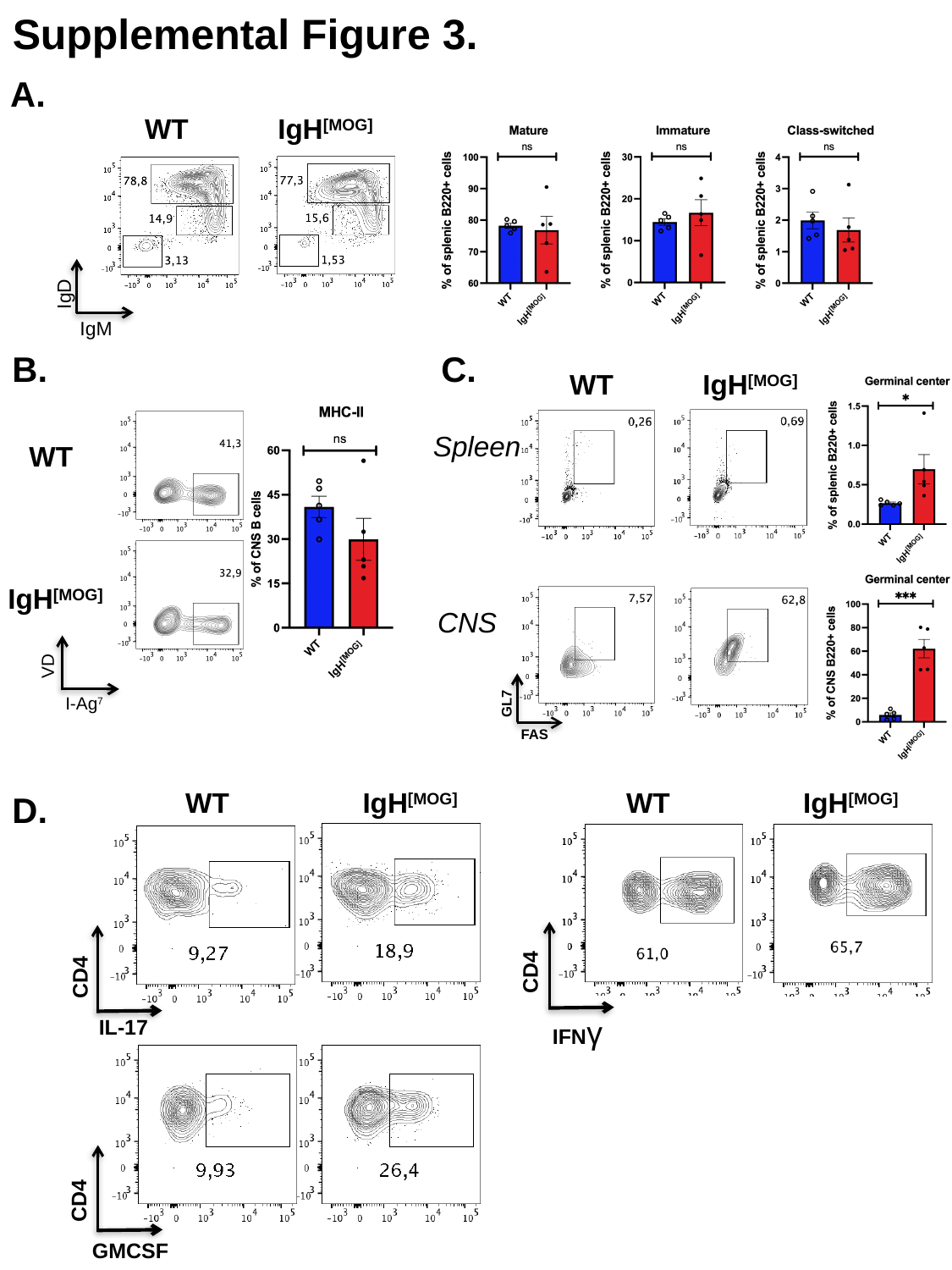

Supplemental Figure 3.
A.
WT
IgH^[MOG]^
IgD
IgM
B.
C.
WT
IgH^[MOG]^
Spleen
WT
IgH^[MOG]^
CNS
VD
I-Ag^7^
GL7
FAS
WT
IgH^[MOG]^
WT
IgH^[MOG]^
D.
CD4
CD4
IFNγ
IL-17
CD4
GMCSF

### Slide 4
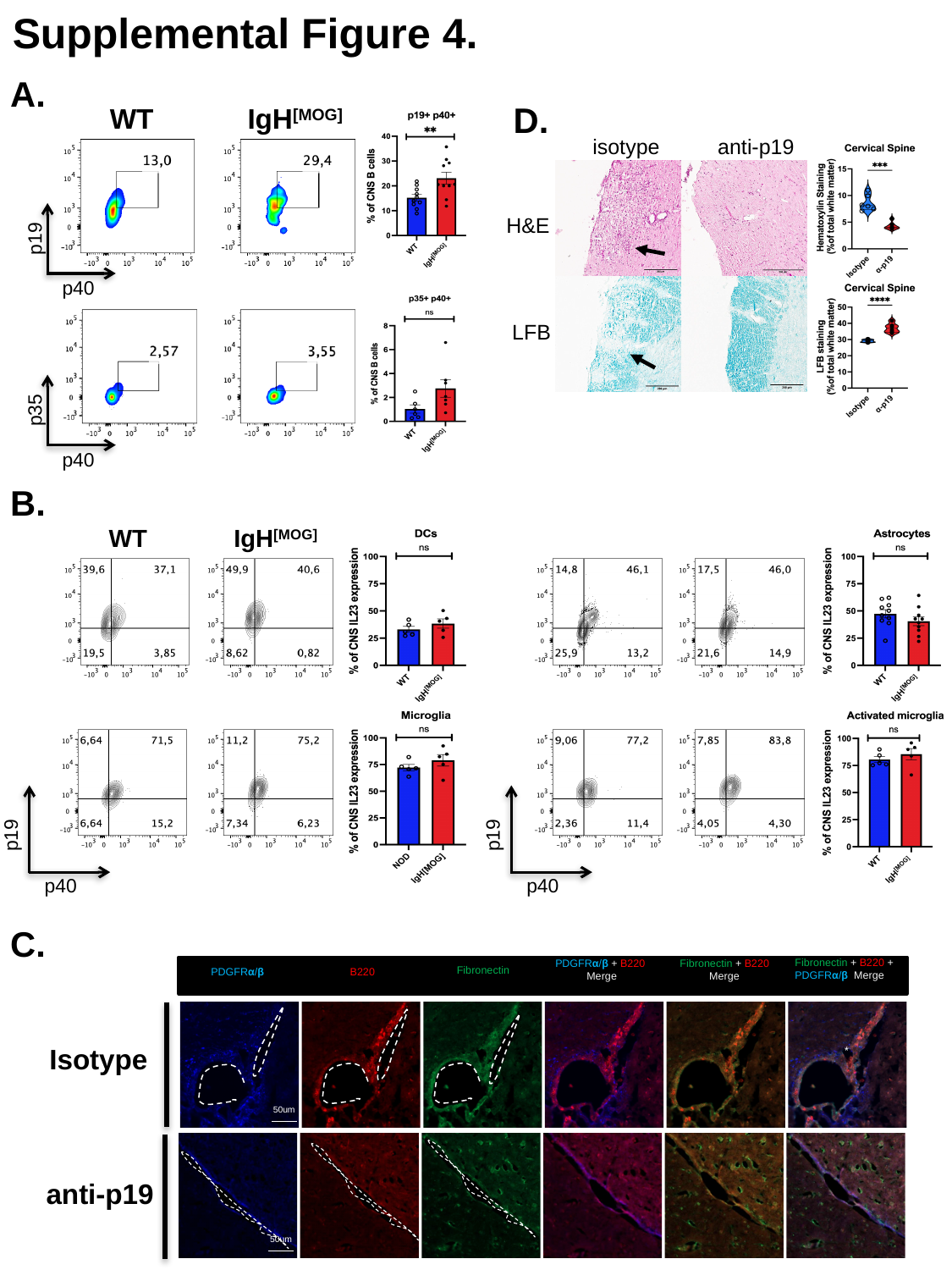

Supplemental Figure 4.
A.
D.
WT
IgH^[MOG]^
p19
p40
p35
p40
anti-p19
isotype
CD4
H&E
LFB
B.
WT
IgH^[MOG]^
p19
p40
p19
p40
C.
PDGFRα/β + B220 Merge
Fibronectin + B220
 Merge
Fibronectin + B220 + PDGFRα/β  Merge
Fibronectin
PDGFRα/β
B220
Isotype
50um
*
anti-p19
50um

### Slide 5
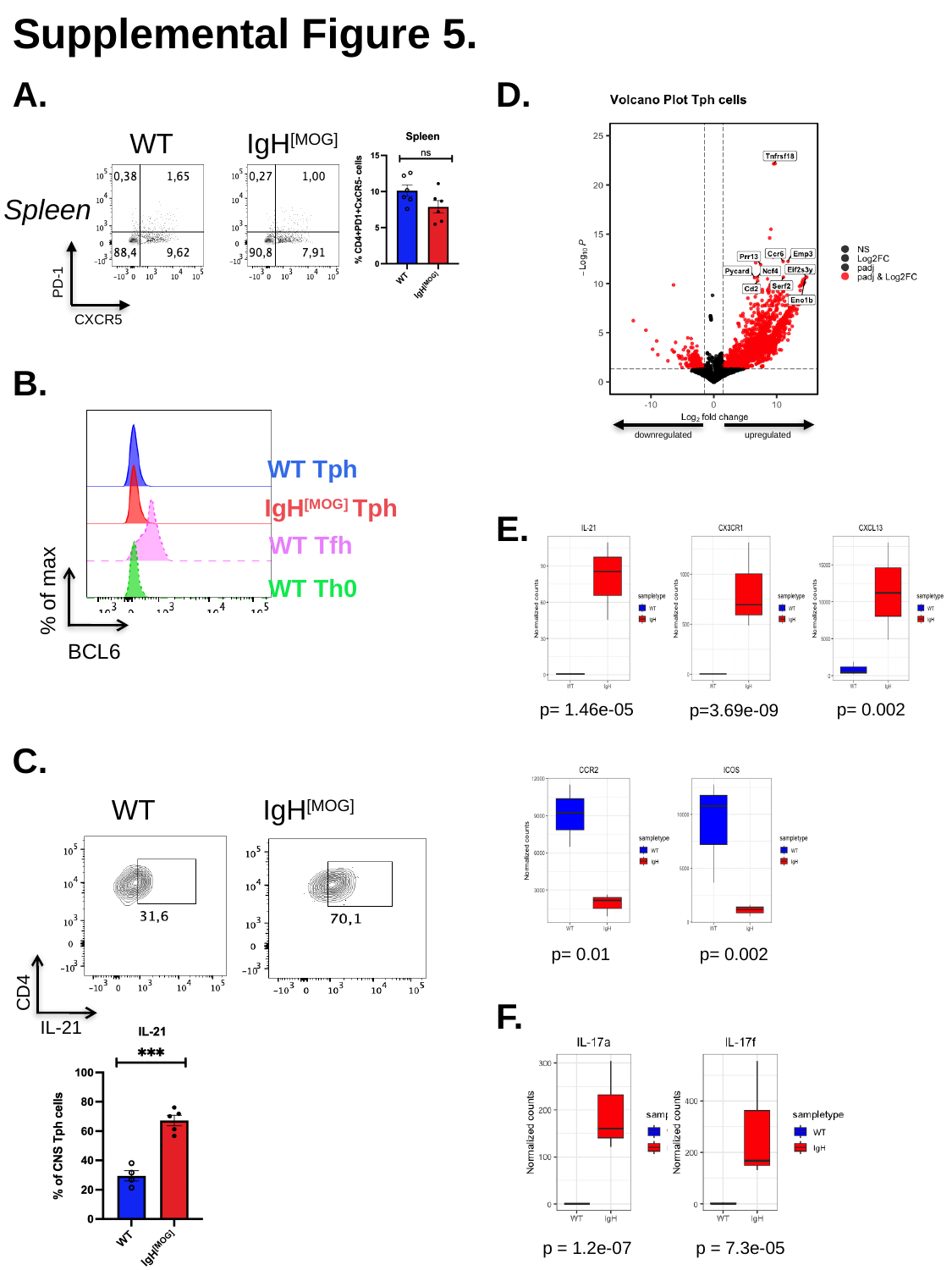

Supplemental Figure 5.
A.
D.
downregulated
upregulated
WT
IgH^[MOG]^
Spleen
PD-1
CXCR5
B.
WT Tph
IgH^[MOG]^ Tph
WT Tfh
WT Th0
E.
% of max
BCL6
p= 1.46e-05
p= 0.002
p=3.69e-09
C.
IgH^[MOG]^
WT
CD4
IL-21
p= 0.01
p= 0.002
F.
p = 1.2e-07
p = 7.3e-05
